# Temporal variation in the behaviour of a cooperatively breeding bird, Jungle Babbler (*Argya striata*) at diel and seasonal scale

**DOI:** 10.1101/2021.11.06.467590

**Authors:** Soniya Devi Yambem, Manjari Jain

## Abstract

Time is an important and limited resource that can drive the trade-off between various essential activities in the lives of animals. Group-living animals need to perform different behaviour to meet their individual needs and also participate in group activities. They must, therefore, partition the available time between these activities which may vary considerably with environmental and ecological conditions. We examined time-activity budget of a cooperative passerine, Jungle Babbler (*Argya striata*) and how their behaviour vary across diel and seasonal scales. A repertoire of 13 behaviour was recorded of which 12 behaviour that occur throughout the year were examined further in detail. This included individual behaviour such as foraging, grooming, rest, shower and group behaviour such as allogrooming, movement, play, sentinel, mobbing and inter-group fight. Our results indicate that most of the time (about 70%) was spent performing individual behaviour and the remaining time (about 30%) was allocated to social behaviour. We also found almost all behaviour varied across diel and seasonal scale with respect to proportion of time spent performing them. This highlights the impact of environmental factors on how animals partition their time to perform various activities. Our study also lays the foundation for future studies examining the role of ecological factors such as habitat type and predation pressure in driving these patterns of behaviour in Jungle Babblers.

## Introduction

Animals carry out different behaviour on a daily basis throughout their lifetime in order to survive and reproduce. These behaviour are allocated differential amount of time which is called as ‘time-activity budget’. Allocation of time to different activities is dependent on the metabolism, energetic constraints as well as the importance of behaviour in the sustenance of life of the animal (Halle and Stenseth 2000). Given that time itself is a limited resource, there is often a trade-off between the allocation of time to different behaviour which may depend on the physical state of the animal, environmental and ecological factors (Pollard and Blumstein 2008). For instance, foraging behaviour is crucial for the survival of the animal and helps in acquiring energy to perform other activities. It is, therefore, expected to have a larger proportion of time allocated to it leaving less time for other activities such as resting or grooming (Kramer 2001). However, in social animals, allocation of time become more complex since the animals not only need to devote enough time to successfully gather resources and reproduce but also to spend time on activities that help in maintaining social bonds such as allogrooming (Boccia et al. 1982; Dunbar 1991) and play (Pozis-Francois et al. 2004) and in those that aid in group coordination such as movement or sentinel behaviour (Hollén et al. 2008). Activities involved in maintaining or strengthening social bonds correlate with direct or indirect fitness of the animal (Silk 2007; Dunbar et al. 2009) and are vital for social animals. Failure to manage time between the various activities may have serious consequences on the number of calories consumed and exhaustion. In addition, for social animals it may also impact social bonding and thereby group dynamics.

The behavioural states of an animal are grouped into two broad categories: ‘activity’ and ‘rest’. Activity behaviour such as foraging, territory defence, exploration of novel territory and finding mates etc., require disproportionately more energy while rest behaviour such as sleeping/resting, grooming, playing etc. require less energy. Even though activity behaviour is crucial for survival, one cannot continuously remain in a high activity state. Such behaviour are typically alternated by resting behaviour to restore energy. The transition of one behavioural state to another across a 24 h cycle, on a daily basis, leads to the development of a temporal pattern which we generally called as diel activity pattern or activity pattern of a species (Halle and Stenseth 2000). Based on the activity pattern in different times of the day, animals are categorized as nocturnal (active at night), diurnal (active during daytime), crepuscular (active at twilight) and cathemeral (active almost equal proportion of time during day and night) (Ikeda et al. 2016). This variation in diel activity patterns across species creates a behavioural niche, allowing the coexistence of different species (Monterroso et al. 2014; Sunarto et al. 2015).

Activity patterns can vary within a species, driven both by abiotic and biotic factors. Biotic factors that may affect the activity pattern in animals include the activity patterns of other species including predators (Lima and Dill 1990), humans (Banerjee and Bhadra 2021) and competitors (Blanchet et al. 2008). Among abiotic factors, the course of seasons and the time of the day are crucial as they significantly impact changes in various environmental parameters such as light intensity, temperatures (Steiger et al. 2013). For instance, dark-eyed juncos initiate feeding early in dim light to replenish their low reserved energy and terminate their activity at high light intensity before the risk of predation increases (Lima 1988). European ground squirrels spends more time resting when the ambient temperature is high during midday (Váczi et al. 2006). Emergence and roosting behaviour, in particular, are likely to be strongly influenced by ambient light conditions and this is likely to change across different seasons. Understanding the activity-patterns and the extent to which these change with various factors provides insights into the ecology of animals. In social vertebrates, most studies on activity budget and the temporal variation in activity patterns have been carried out on primates (Rasmussen 1985; Isbell and Young 1993; Zhou et al. 2007; Back et al. 2019; Li et al. 2019). While many avian species are social, yet, to our knowledge, similar studies are lacking in social passerines.

Jungle Babbler (*Argya striata*) is a cooperatively breeding bird belonging to the order passeriformes and family leiothrichidae (Cai et al. 2019). They are found throughout lowland India, mainly in tropical woodland and scrub vegetation with a preference for human neighbourhood (Andrews and Naik 1970; Gaston 1977). They live in groups of 3-20 individuals that engage in many social behaviour and maintain territory ranges from 0.0014 to 0.016 km^2^ (Andrews and Naik 1970). They possess a complex vocal repertoire of calls that mediate various social behaviour such as foraging, movement, sentinel activity and brood care (Yambem et al. 2021). In this study we aimed to address the following question 1) Do Jungle Babblers differentially allocate time to different behaviour? 2) Does the time spent performing different behaviour changes across diel and seasonal scale? 3) Does roosting and emergence from roost vary with ambient light conditions and seasons? We hypothesised that the birds would allocate higher proportion of time towards sustenance as compared to those involved in social bonding. Towards this our prediction was that the highest proportion of their time budget would be allocated to foraging as it is an important behaviour that determines sustenance. We also hypothesized that the fraction of time allocated for behaviour is likely to be influenced by environmental features. Towards this we predicted that the activity patterns will vary across the different seasons due to changes in ambient temperature and foliage cover across seasons. Following our hypothesis, we also expected both roosting and emergence to vary in relation to ambient light conditions and seasons. Additionally, we examined sentinel duty in detail to understand what proportion of foraging activity had a sentinel on duty and whether this varied with season. Gaston (1977) examined sentinel behaviour in Jungle Babblers during winter season but did not examine seasonal variation in this behaviour.

According to Enright (1970), “No description of where an animal lives and what it does can be complete without considering when the activity takes place because animals are obviously adapted to perform given activities at given environmental times: certain seasons, times of day, or phases of the tides”. This study will provide insights into the ecology of a tropical social passerine. It will also provide novel data on the impact of environmental condition on activity patterns of a social passerine, thereby furthering our understanding of factors that determine trade-off in time investment across different behaviour in social animals. Such data are lacking in social avian species and specifically lacking from the tropics that are home to many social animals.

### Materials and methods Study species and study site

The study was conducted in Mohali region, located in the eastern part of the Punjab state in India (30°36’ and 30°45’N latitude and 76°38’and 76°46’E longitude), which covers an area of about 116.50 Km^2^ (Tur et al. 2011). The climate of Mohali comes under ‘Cwa’ category. Mohali has a humid subtropical climate with dry winter, hot summer, humid monsoon and a short transitional period of postmonsoon (Kottek et al. 2006). For ease of observation and logistical considerations, all observations were made on institute campuses where a healthy population of Jungle Babblers is known to exist and have been monitored regularly over several years. A total of three locations were selected across two institute campuses: 2 on IISER Mohali campus and 1 on NIPER campus, such that each site was at least 500 m away. The campuses have a mix of gardens, open grassland, plantation and natural closed-canopy woodland. The area was dominated by deciduous plant species as well as weedy species. The plants species include *Populus deltoides, Bombax ceiba, Bauhinia purpurea, Schleichera oleosa, Dalbergia sissoo, Ficus religiosa, F. glomerata, F. virens, Vachellia nilotica, Pongamia pinnata, Morus alba, M. nigra, Psidium guajava, Leucaena leucocephala, Chukrasia tabularis, Callistemon* sp., *Lantana camara, Ricinus communis* and *Cannabis* sp.

### Data collection

All the observations were done using 10 × 50 binoculars (Nikon Monarch) at a distance of >5m. *Ad libitum* sampling (Altmann 1974) for 5 months was carried out so that the animals can be habituated to the observers’ presence and the observers can understand the patterns of movements, rough territory size and list of observable behaviour. For this study, exact group identity was not required since we aimed to examine activity pattern and diel and seasonal variation in different behaviour exhibited by the species. Nonetheless, since Jungle Babblers are known to be territorial (Andrew and Naik 1970), we could ensure that we were sampling distinct groups by sampling in distinct localities far away from each other. However, sometimes different groups came together towards the same foraging ground (Andrew and Naik, 1970) but since we did not aim to examine group-level differences in these behaviour, it did not pose as a limitation. The repertoire of 13 different behaviour observed in the focal species are summarised for reference in Table 1. Some of the common behaviour are shown in pictures (Fig. 1). This list of behaviour are based on our own observations which is validated by previous studies on Jungle Babblers (Andrew and Naik 1970; Gaston 1977 and Yambem et al. 2021).

**Table 1.** Ethogram of Jungle Babbler with 13 behavioural repertoires. Shaded rows represents the behaviour which were used in this study for time-activity budget and activity pattern. * indicates the behaviour which were grouped into ‘other’ behaviour and + represents those behaviour which are associated with vocalization (Yambem et al. 2021)

| Behaviour | Description |
| --- | --- |
| 1. Foraging + | Process of finding and obtaining food by hopping and pecking on the leaf litter, foliage, bark crevices and ground. |
| 2. Grooming | Activity of cleaning or maintaining one’s own body with its beak. |
| 3. Rest | One or more individuals perch on a branch next to each other with eyes open or closed, not partaking in other activities. |
| 4. Shower * | Quickly dipping body in a shallow pool of water and subsequently shaking off water from feathers. |
| 5. Allogrooming | Two or more individuals groom each other by pecking lightly. |
| 6. Movement + | Flying as a group, with or without vocalizations, from one spot to another within a distance of 10-20 m by making quick stops in between. |
| 7. Play * | “Two or more birds engaged in a mock fight in which some lie on the ground more or less passively, while others rolled on top of them, or pecked them deliberately but gently” or sometimes chase one another in the air just above the ground or between the trees (Gaston 1977). |
| 8. Sentinel + | One, sometimes two, individuals perch on an elevated platform, while other group members forage on ground. |
| 9. Mobbing * + | Attacking or chasing the predator, by some individuals or whole group members. |
| 10. Inter-group<br>fight * + | Aggressive interactions between members of two groups that encounter each other. Interactions mainly mediated through vocalizations and sometimes followed by physical combat. |
| 11. Emergence | All members of a group emerge from the roost one by one. |
| 12. Roosting | Group members perched on one or two branches in a clumped manner, either facing in the same or in opposite direction. |
| 13. Parental care<br>+ | Involves the activity of feeding, grooming and guarding the nestling/fledgling, or prompting the fledglings to fly. |

**Fig. 1.**
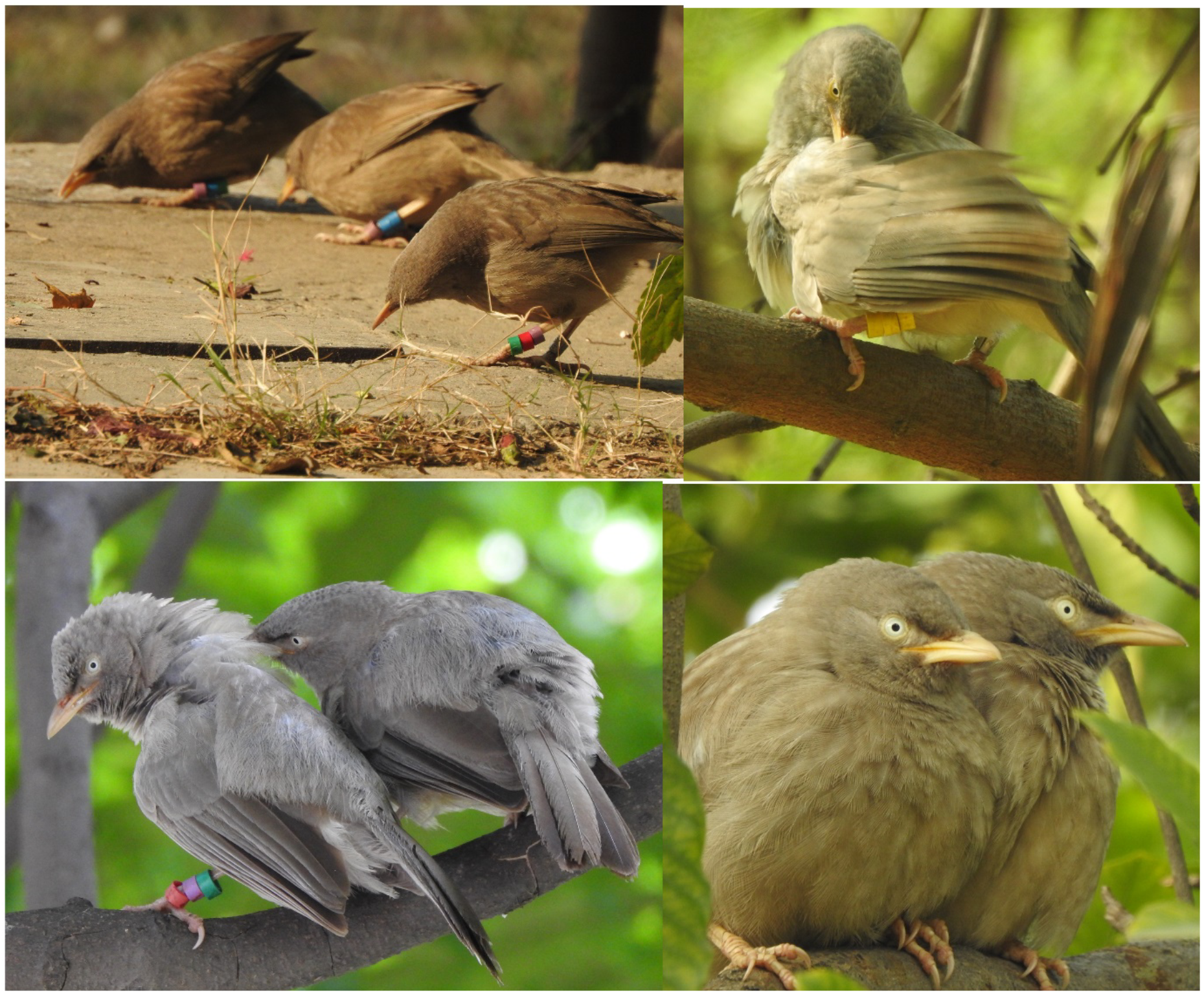
Jungle Babbler showing various behaviour. From top left to bottom right: foraging, grooming, allogrooming and rest

The behavioural data for this study was collected between October 2016 to September 2017. Of the 13 behaviour listed, 12 occurred throughout the year. Parental care was not included in this study as it is limited to breeding time. To record the activity pattern between 5:00 to 20:00, 6 hours of observations were carried out in 2 sessions in a day with a break of at least 3 hours in between. Each session included 3 sampling hours of observations and a sampling hour was divided into a sequence of alternating 5-minute periods of observation and rest each in order to avoid exhaustion of the observer. Timing of the observations was alternated across days such that all time slots between 5:00 to 20:00 were covered at least once in a week. Every 1 hour time slot was sampled at least on 20 days, spread across the year, in the study period (Table S1). However, due to adverse weather conditions and poor visibility, only 9 days of sampling was carried out in the month of January (Table S2). The frequency of visiting the three different locations was kept the same. The sampling technique used was instantaneous scan sampling (Altmann, 1974) in which all the behavioural activity performed by observable group members as well as the duration of the activity was noted using a digital stopwatch (Marathon Adanac 3000). If all the birds being observed went out of sight before completing the 5 minutes, then the observation was truncated and data from that sampling period was discarded in order to maintain uniformity across sampling times.

Observations on emergence from and return to the roost sites of Jungle Babblers were done one hour before predicted sunrise (5:19-7:21) and sunset (17:32-19:30) times, respectively. The time of the return to roost and emergence was noted down only when all the group members settled down and clumped together on a branch of a tree and when all the group members flew out from the roosting tree respectively. The light intensity at the time of roosting and emergence was also noted down using light meter (Lutron Lx 1102).

### Data analysis

The time activity budget was calculated for 10 out of the 13 observed behaviour (highlighted in colour in the Table 1). Emergence and roosting behaviour were analysed separately since emergence is an event behaviour and roosting occurred at the end of the day and through night time. The time activity-budget was calculated as the proportion of time spent performing each of the behaviours by dividing the number of scans in which a particular behaviour was exhibited by the total number of scans summed across all behaviour. This was then converted into percentage values representing the percentage of time spent for each of the different behaviour. In addition, we calculated the proportion of time spent for each behaviour in each of the 15 sampling hours (5:00 – 20:00), averaged across all months, to examine diel pattern of the behaviour and for each month of the year, averaged across all sampling hours, in order to examine seasonal variation in activity pattern. To compare time spent on individual versus social behaviour, the 10 behaviour were grouped into two categories: individual and social behaviour. Those behaviour that do not require participation of other members of the group were designated as individual behaviour (foraging, grooming, rest and shower), whereas those that require participation of other in the group were labelled as social behaviour (allogrooming, movement, play, sentinel, mobbing and inter-group fight). For ease of analyses, rarely observed behaviour (that occurred in < 1% of all observations) were grouped into a single behavioural category called ‘other’ (Table 1). To examine seasonal variation in activity patterns, data were arranged into 4 seasons: winter (December to February), summer (March to June), monsoon (July to September) and postmonsoon (November to December). This grouping resulted in fewer days in some seasons and more in others, however, it represents the most relevant categorization given the meteorological conditions experienced in Mohali. Since the data for seasonal variation were calculated as averages, this inequality should not matter.

### Statistical analyses

All the statistical analyses were carried out in R version 4.1.0 (R Core Team, 2021). To check the influence of factors such as time of the day (diel pattern) and length of the day on different behavioural activities, Generalised Linear Model (GLM) was run for each behavioural activity. Response variable for the model was the proportion of time allocated to each behaviour: foraging, grooming, rest, allogrooming, movement, sentinel and ‘other’. Since the response variable was the proportion data, family “quassibinomial” with link “logit” was applied in the model. To examine seasonal variation in the proportion of time spent on behavioural activities as well as on the timing of emergence and roosting, Kruskall Wallis test was run. Mann-Whitney U test was further carried out to make pairwise comparisons between seasons. To examine the difference between the light intensity at the time of emergence and roosting, Mann-Whitney U test was carried out.

## Results

### Time activity budget

In one year, a total of 18,178 behavioural records (pooled across all behaviour) across 12, 330 scan samples collected across 192 days of observation were obtained by combining all the observation from the three locations. Detailed sample size for every sampling hour and month are given in Table S1 and S2. Time-activity budget are represented in percentage in Fig. 2. Our findings suggested that Jungle Babblers spent around 69% of time on individual behaviour and the remaining 31% was allocated to social behaviour (Fig. 2). Individual behaviour such as foraging, grooming, rest and shower are necessary for sustenance and maintenance, whereas, social behaviour such as allogrooming, movement, play, sentinel, mobbing and inter-group fight are required to maintain group stability. Amongst different behaviour, the highest amount of time was devoted to foraging (56.6%) followed by sentinel (16.65%). Further, we found that that on an average, a sentinel was present only about 32% of the time while Jungle Babblers foraged. The remaining time, no sentinel was found to be on duty. Further, we found that Jungle Babblers spent almost equal amount of time (χ^2^ = 0.047, df =1, p = 0.82) on grooming (10.17%) and allogrooming (9.21%) and the least amount of time was devoted to movement (3.45%) and resting (1.61%). The remaining amount of time was allocated to several behaviour that were pooled as ‘other’ (1.79%) (Fig. 2).

**Fig. 2.**
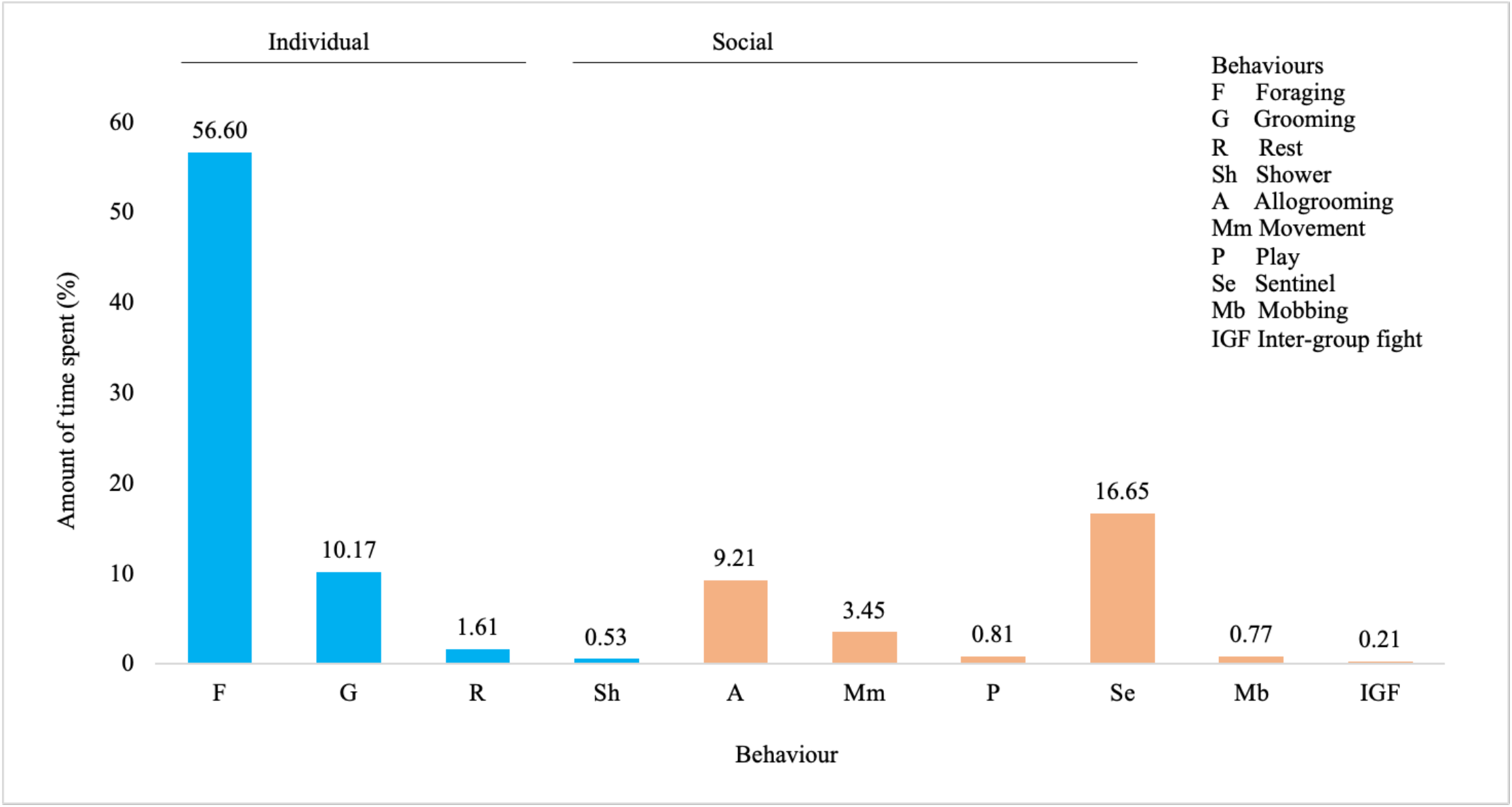
Amount of time spent (represented in percentage) on each behavioural activity calculated from 18,178 total scans of all behaviour collected from 192 days between October 2016 and September 2017

### Diurnal and seasonal variation in activity pattern

The outputs of GLM results showed that foraging, grooming, rest, sentinel and movement behaviour exhibited diurnal activity pattern whereas, allogrooming and ‘other’ behaviour did not show any diurnal pattern (Fig. 3 and Table 2). Sentinel, rest and movement behaviour increased with the time of the day (Table 2) while foraging and grooming decreased with the time of the day (Table 2). Diurnal activity pattern of allogrooming, rest and sentinel varied with the length of the day wherein, allogrooming and rest increased but sentinel activity decreased with the length of the day (Table 2). Results of Kruskal-Wallis test showed that all behaviour varied across seasons (Fig. 4 and Table 3a). The proportion of time spent foraging was highest during postmonsoon and lowest during winter (Fig. 4). Pairwise comparisons across seasons revealed significant difference in time spent foraging across most seasons (Table 3b). Grooming and allogrooming were high during monsoon and both rest and movement activities were found to be high during summer and ‘other’ activity during monsoon (Fig. 4). While Jungle Babbler foraged, the percentage of time that a sentinel was present during foraging activity varied significantly across seasons wherein, in winters a sentinel was found to be on duty 48% of time but only 14% of time during the monsoon (Fig. 5 and Table 3c).

**Table 2.** Summary of GLM results examining the effect of time and length of the day on the proportion of time spent on different behaviour

| **Behaviour** | *Predictor* | *Estimates* | *Std.error* | *p* |
| --- | --- | --- | --- | --- |
| **Foraging** | Intercept | 0.40 | 0.33 | 0.23 |
|  | Time | -0.03 | 0.01 | <0.0001 |
|  | Length | 0.03 | 0.03 | 0.32 |
| **Grooming** | Intercept | -1.34 | 0.58 | 0.02 |
|  | Time | -0.09 | 0.02 | <0.001 |
|  | Length | 0.02 | 0.05 | 0.73 |
| **Rest** | Intercept | -10.26 | 1.73 | <0.0001 |
|  | Time | 0.01 | 0.04 | 0.01 |
|  | Length | 0.38 | 0.13 | <0.001 |
| **Allogrooming** | Intercept | -4.97 | 0.61 | <0.0001 |
|  | Time | 0.02 | 0.01 | 0.259 |
|  | Length | 0.19 | 0.05 | <0.001 |
| **Sentinel** | Intercept | 0.84 | 0.42 | 0.04 |
|  | Time | 0.05 | 0.01 | <0.0001 |
|  | Length | -0.26 | 0.03 | <0.0001 |
| **Travel** | Intercept | -5.76 | 1.17 | <0.001 |
|  | Time | 0.20 | 0.03 | <0.001 |
|  | Length | -0.001 | 0.09 | 0.99 |
| **Other** | Intercept | -6.43 | 1.31 | <0.001 |
|  | Time | 0.03 | 0.03 | 0.33 |
|  | Length | 0.19 | 0.12 | 0.06 |

**Table 3.** Summary of a) Kruskal-Wallis ANOVA examining effect of season on the proportion of time spent on different behaviour and the timing of emergence and roosting b) Mann-Whitney U test (p values) for pairwise comparison between different seasons for proportion of time spent on different behaviour and the timing of emergence and roosting c) Mann-Whitney U test (p values) for pairwise comparison between different seasons for proportion of time spent on sentinel duty while foraging.

| Behaviour | $\chi^2$ | df | p |
| --- | --- | --- | --- |
| Foraging | 14.09 | 3 | <0.01 |
| Grooming | 9.47 | 3 | 0.02 |
| Rest | 11.79 | 3 | <0.01 |
| Allogrooming | 8.45 | 3 | 0.03 |
| Movement | 21.32 | 3 | <0.001 |
| Sentinel | 93.06 | 3 | <0.001 |
| Other | 7.96 | 3 | 0.04 |
| Emergence | 22.33 | 3 | <0.001 |
| Roosting | 14.71 | 3 | <0.01 |

9 b)
| Seasons | Foraging | Grooming | Rest | Allogrooming | Movement | Sentinel | Other | Emergence | Roosting |
| --- | --- | --- | --- | --- | --- | --- | --- | --- | --- |
| Postmonsoon-Winter | **<0.01** | 0.46 | 0.61 | 0.40 | 0.65 | **<0.01** | 0.91 | 0.12 | 0.26 |
| Postmonsoon-Summer | **<0.01** | 0.99 | 0.057 | 0.40 | **<0.01** | 0.71 | 0.87 | **0.027** | 0.09 |
| Postmonsoon-Monsoon | 0.40 | **0.026** | 0.99 | 0.07 | **<0.001** | **<0.001** | 0.06 | 0.64 | **<0.001** |
| Winter-Summer | 0.31 | 0.4 | **0.011** | 0.06 | **0.034** | **<0.001** | 0.77 | **<0.001** | **0.023** |
| Winter-Monsoon | **0.01** | 0.12 | 0.59 | **<0.01** | 0.09 | **<0.0001** | **0.038** | 0.39 | 0.05 |
| Summer-Monsoon | **0.034** | **<0.01** | **0.012** | 0.21 | **<0.0001** | **<0.0001** | **0.022** | **<0.001** | **<0.01** |

6 c)
| Seasons | p |
| --- | --- |
| Postmonsoon-Winter | <0.001 |
| Postmonsoon-Summer | 0.45 |
| Postmonsoon-Monsoon | <0.0001 |
| Winter-Summer | <0.001 |
| Winter-Monsoon | <0.001 |
| Summer-Monsoon | <0.0001 |

**Fig. 3.**
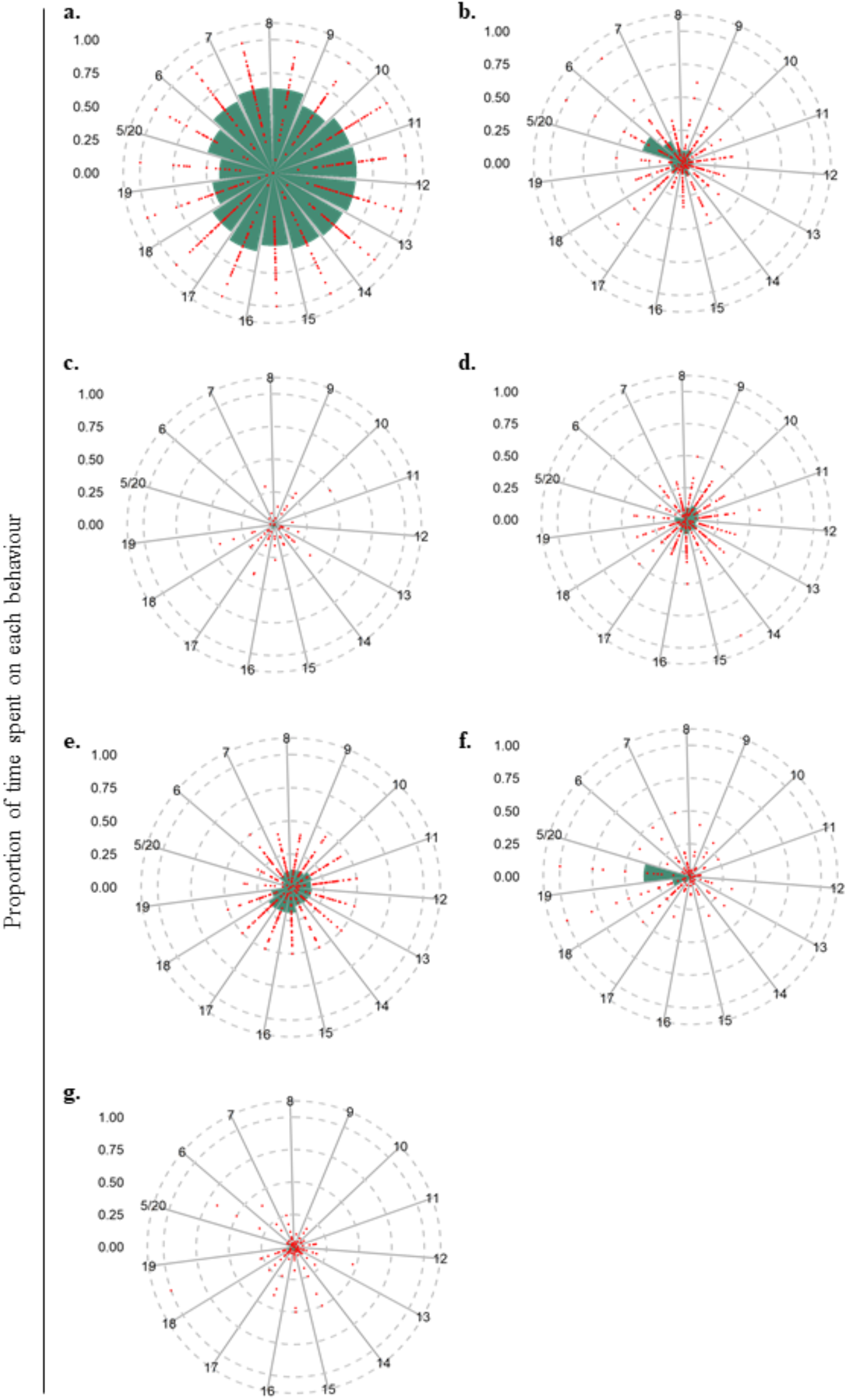
Polar plot showing the proportion of time spent on each behavioural activity across 15 hours (5:00-20:00): a) foraging, b) grooming, c) rest, d) allogrooming, e) sentinel, f) movement and g) ‘other’, at different time of the day. Each point represents the proportion of time spent on each behaviour in each sampling hour of a day and bar represents the average proportion of time spent on each behaviour in each sampling hour across one year.

**Fig. 4.**
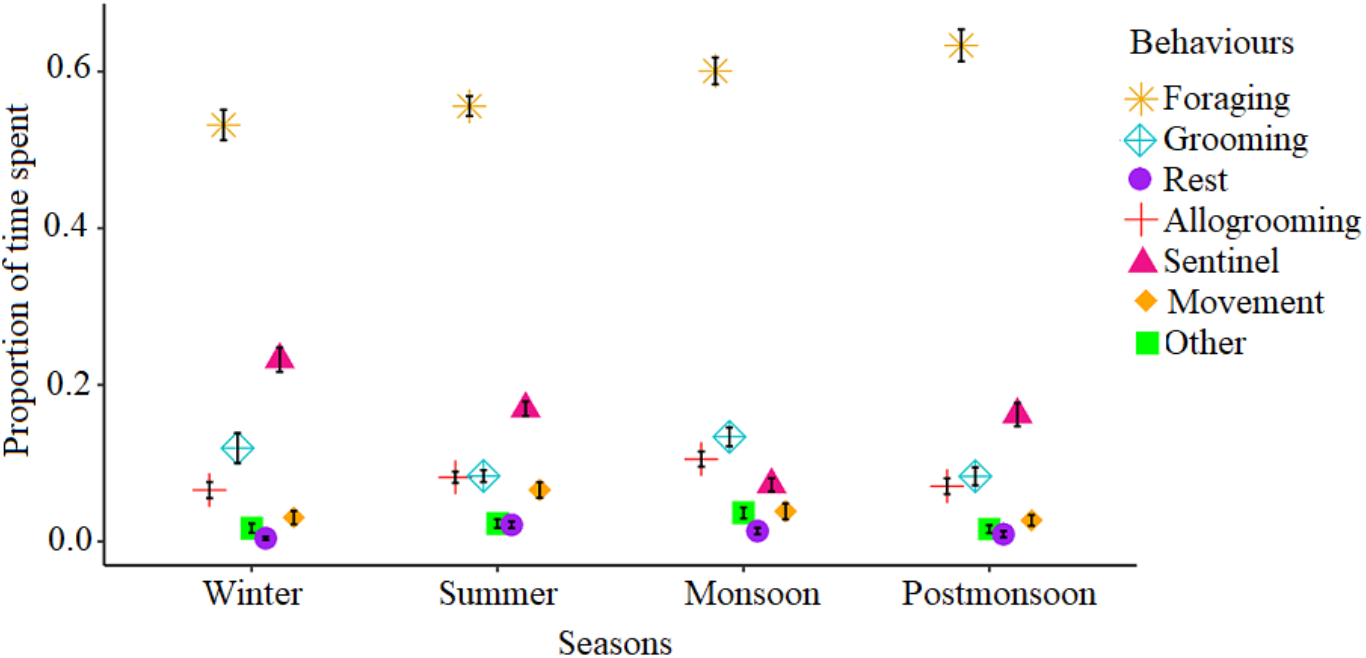
Mean ± SE plot for the proportion of time spent on each behavioural activity across seasons.

**Fig. 5.**
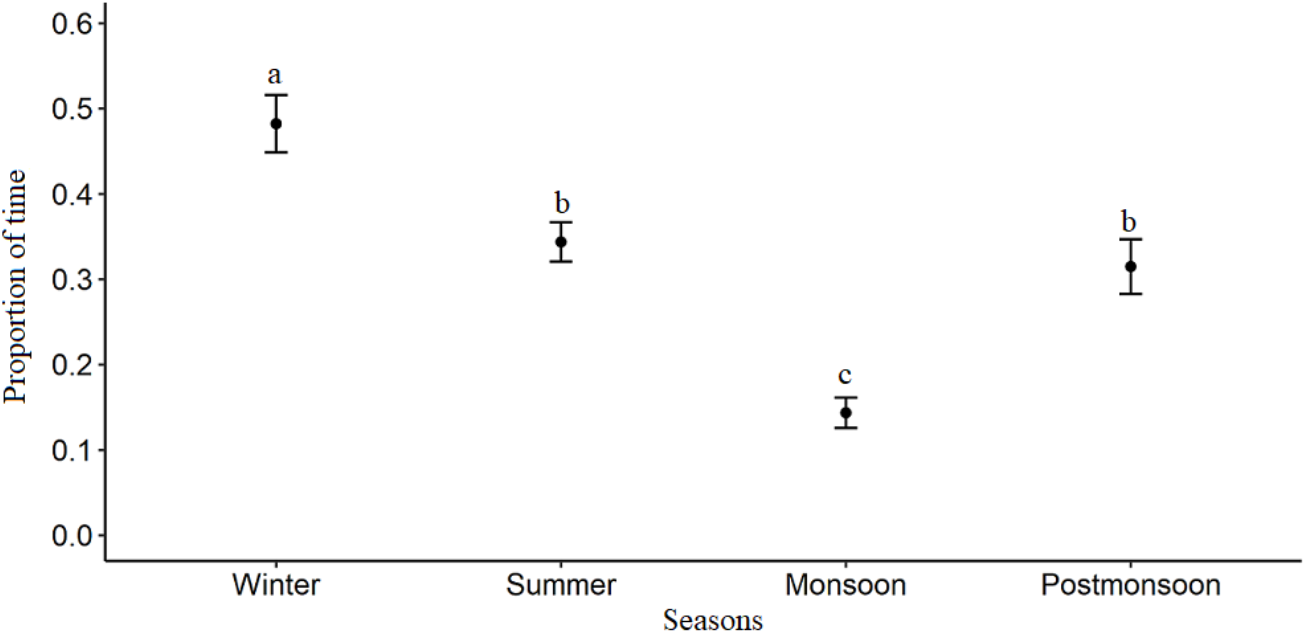
Mean ± SE plot for the proportion of time, a sentinel was present while the group members were foraging on the ground. Different letters indicate significant difference with p < 0.05

### Emergence and roosting

Results of Kruskal-Wallis test showed that the timing of emergence and roosting varied with season (Fig. 6a and Table 3a.). Pairwise comparisons showed that Jungle Babblers emerged from their roosts earliest during summer (Fig. 6a and Table 3b). Pairwise comparisons also revealed that Jungle Babblers returned to their roosts significantly earlier during winter and monsoon, which did not differ significantly from each other (Fig. 6a and Table 3b). Light intensity at the time of roosting was significantly higher than the emergence (Mann-Whitney U: p <0.001, Fig. 6b).

**Fig. 6.**
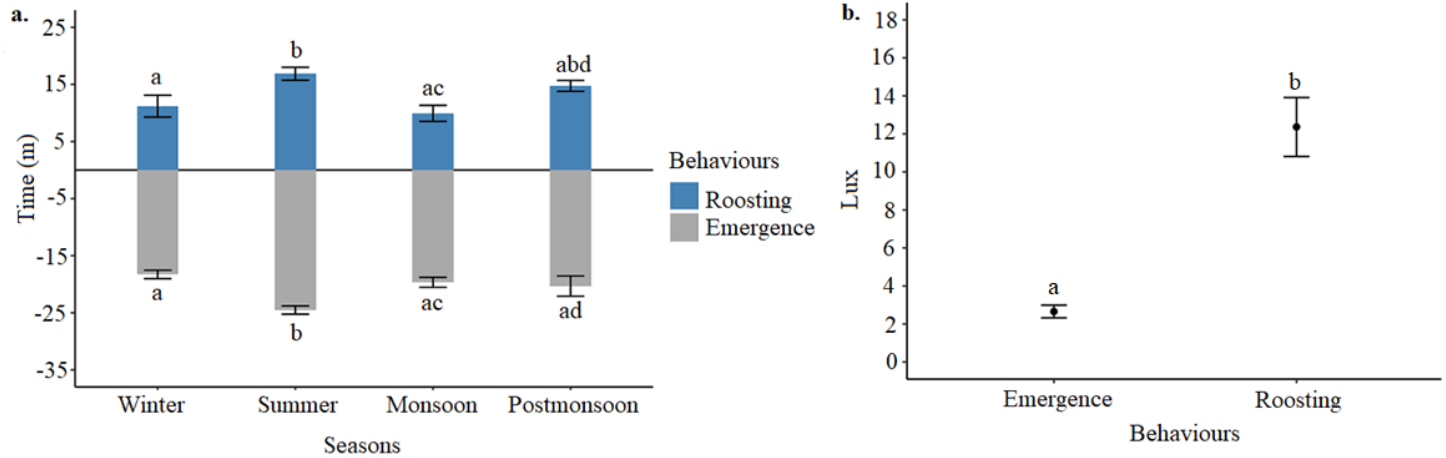
a) Emergence and roosting time (mean ± SE) in relation to sunrise and sunset respectively during different seasons. The number of days of observation of emergence and roosting in winter, summer, monsoon and postmonsoon are 25, 17, 15, 12 and 26, 23, 15, 21 respectively. b) Light intensity in lux unit (mean ± SE) at the time of emergence and roosting. The number of days of observation (light intensity) for emergence and roosting are 50 and 53 respectively. Different letters indicate significant difference with p < 0.05

## Discussion

### Activity budget

Time is an important resource and its availability to animals is limited. Examining how animals partition their time between various activities can reveal much about their ecology. Gadagkar and Joshi (1983) in their study on the social wasps, *Ropalidia marginata* used time-activity budget data to reveal behavioural castes in the species that lacks morphological differences between individuals, unlike other social insects. Dunbar (1992) showed that spending a lot of time on activities that are simply required for sustenance come at the cost of time spent for social behaviour and may result in group instability and thereby limit group size. This makes the studies on time activity budgets of social animals essential to understand the evolution of sociality. We found that Jungle Babblers devoted around 70% of their time on individual behaviour which are required for one’s sustenance and maintenance and the remaining 30% for social behaviour. The relatively high proportion of time spent on individual behaviour is mainly dominated by foraging behaviour. This is unsurprising, given that foraging helps in gaining energy, which is required for survival, growth as well as reproduction (Kramer, 2001). Similar findings have been reported in urban Capuchin monkeys (80%) and Shelducks (60%) in which most of their time is allocated in foraging (Back et al. 2019; Bensizerara and Chenchouni 2019). However, Vervet monkeys have been reported to spend much less time in foraging (30-40%, Isbell and Young 1993) in comparison.

Grooming and shower behaviour come under the same category of avian maintenance which involves removing dirt, parasite and maintaining hygiene (Clayton and Cotgreave 1994). Clayton and Cotgreave (1994) stated that grooming is a time consuming activity and in a comparative study across 62 species of birds, they showed that on an average 9.2% of time was devoted to grooming which is similar to the amount of time Jungle Babblers allocated to grooming (10.17%). Jungle babblers spent only a small percent of the time in resting (1.61%) during the day. However, it must be noted that the time between roosting and emergence is devoted exclusively to resting and has not been included in the analyses. Further, during the day time, the time spent grooming may also serve similar function as rest behaviour, providing a break from high activity and thereby avoiding exhaustion. Resting is important for the physiological processes such as digestion and thermoregulation, however, the time spent during resting is considered to be free and can be utilized for other activities when required (Herbers 1981; Dunbar 1992). Thus, in social animals, it is expected that most of the resting time will be devoted to social activities (Dunbar and Dunbar 1988). In fact, Jungle Babbler also devote nearly 10% of their time in allogrooming, which too is a less energy intensive activity but is crucial for social bonding. Besides maintaining social bond (Isbell and Young 1993; Cox 2012; Picard et al. 2020), allogrooming also serves a similar role as grooming behaviour in removing dirt, parasites and maintaining hygiene (Sparks 1967; Sachs 1988) suggesting that both the behaviour share somewhat similar kind of ecological function. In fact, our finding suggests that the proportion of time dedicated to grooming and allogrooming are almost equal.

Among all social behaviour in Jungle Babblers, most of the time was allocated to sentinel behaviour (16.65%, averaged across all months of the year) with respect to time-activity budget. This activity was carried out by 1 individual and rarely by 2 individuals by taking turns. Sentinel activity in Jungle Babblers is associated with a soft vocalization (Gaston 1977; Yambem et al. 2021). Even though there is no visible response from group members towards this vocalization (Yambem et al. 2021), Wickler (1985) suggested that this vocalization might mediate the coordination between the foragers and sentinel in Jungle Babblers. Further, during foraging bouts, we found a sentinel to be present about 32% of the time on an average. This is similar to the findings in Pied Babblers, where too sentinel was found to be present only 30% of the time during foraging bouts (Hollén et al. 2008). Play behaviour has been proposed to help in maintaining social bond, yet Jungle Babblers allocated only 0.81% of their time to play. Since play is physically and socially demanding behaviour (Pozis-Francois 2004), we speculated that for maintaining social bond Jungle Babblers spent more time in allogrooming which is less energy demanding. In fact, studies in chimpanzees have shown that the time allocated to play decreases when allogrooming time increases (Lawick-Goodall 1968). Our finding that Jungle Babblers devoted only 3.45% of time on movement can be partly explained by the fact that we only included flight as part of movement. Locomotion would include walking as well but we were particularly interested in displacement behaviour and not locomotion per se, which is included in foraging implicitly. Further, Jungle Babblers typically have small territory sizes (Andrews and Naik 1970) which limits the opportunity of movement and corroborates our findings. Four behaviour (shower, play, mobbing and inter-group fight) were grouped as ‘other’, yet, put together they consumed only 2.32 % of total time activity budget. This might be because shower and play behaviour are functionally similar to grooming and allogrooming, respectively, by virtue of their similar role of maintaining hygiene and social bonding respectively (Lawick-Goodall 1968; Clayton and Cotgreave 1994; Cox 2012). Besides, mobbing and inter-group fight are aggressive behaviour which are likely to be infrequent anyway.

### Diurnal and seasonal activity pattern

The variation in the activity pattern of an animal depends upon many factors including time of the day (Li et al. 2019), season (Ikeda et al. 2016), predation pressure (Lima and Dill 1990) and sociality (Marshall et al. 2012). From our results, it was found that the most of the behaviour of Jungle Babblers showed both diurnal and seasonal variation except for allogrooming and ‘other’ behaviour that did not show diurnal pattern. We found that the time spent on foraging varied with the time of the day and season irrespective of the length of the day. Reyes-Arriagada (2015) examined diel patterns of activity in three species of forest-dwelling passerines and reported that foraging patterns were nonuniform throughout the day and also varied with season and habitat type. In Jungle Babblers, foraging time decreased with the time of the day. Higher foraging activity early in the day ensures rapid energy gain after an extended period of starvation through the night and also reduces the risk of starvation due to lost foraging opportunities later in the day (Bednekoff and Houston1994). With respect to seasonal variation, Jungle Babblers had highest foraging activity during postmonsoon (63%) and lowest during winters (53%). Yet, the proportion of time spent foraging never dropped below 50% at any given time of the year. This is not surprising given that foraging is an essential sustenance activity. Grooming behaviour in Jungle Babblers was highest in the morning and peaked during the monsoon. This is similar to findings in Java monkeys in which grooming was higher during morning as compared to evenings (Troisi and Schino 1986). Further, the presence of ectoparasite load in White Shifakas (lemurs) has been shown to be higher in monsoon which elicit an increase in grooming to maintain self-hygiene (Lewis 2010). Whether parasite load in Jungle Babblers varies with seasons remains to be examined. Allogrooming did not show any pattern across day. However, day length has a significant effect on allogrooming (increases with an increase in day length). This may be because allogrooming mainly aid in social bonding (Picard et al. 2020), can be carried out at any time of the day (Dunbar 1992). Further, allogrooming was found to increase during monsoon which might be explained by its shared ecological role with grooming behaviour in maintaining hygiene. However, this is in contrast to the findings of Gaston (1977), who found that allogrooming in Jungle Babblers increased during the postmonsoon and winter and was lower during summer and monsoon.

Sentinel activity varied through the day, wherein it increased in the evening. Our results are similar to the findings of Gaston (1977) where it was shown that the sentinel behaviour increased with the time of the day, which might be related to achieving satiation later in the day. An experimental study in Arabian babblers showed that well-fed individuals performed more sentinel duty suggesting that sentinel is a state-dependent behaviour (Wright et al. 2001). The proportion of time allocated to sentinel duty varied significantly with season in Jungle Babblers and peaked during the winters. Gaston (1977) remarked based on qualitative observations that sentinel behaviour in Jungle Babbler was highest during winter. We further inspected sentinel duty as a proportion of foraging time during which sentinel was present and how that varied with season. Our results indicate a peak in sentinel duty during winter, with a sentinel being present 48% of foraging time, that drops to 14% during monsoon. This trend is similar to what was found in Florida scrub jays, where sentinel duty during foraging peaked in winters (75% of total foraging time) and dropped during summer to about 33% (Mcgowan and Woolfenden 1989). The differences in sentinel activity can be attributed by various factors such as canopy cover, predation risk and even group size. In our study, the increase in sentinel activity during winters and summer can be attributed to poor canopy cover during this time of the year. The study sites were dominated with deciduous tree that shed their leaves during winter and early summer, thereby possibly increasing predation risk. Given that time must first be allocated for crucial activities like foraging, time for resting is likely to become available only when other activities are fulfilled (Altmann and Muruthi 1988; Dunbar 1992). Our results agree with this prediction and we found that the time spent resting increased with the time of the day as foraging activity decreased. Besides, we also found that resting increased with the day length. Movement behaviour also showed variation with time of the day and season. It increased during the summer which coincides with the starting of breeding season (Andrew and Naik 1970). During this time the birds may spend more time searching for the nest site, building nest etc that may require frequent displacement.

### Emergence and roosting

The timing of emergence and roosting in Jungle Babblers in relation to time of sunrise and sunset respectively, was different in different seasons. Similar findings have also been reported in Indian Myna that showed diurnal and seasonal variation in the emergence and roosting behaviour under the influence of environmental, physiological and behavioural factors (Mahabal and Vaidya 1989). Jungle Babblers emerged earlier and returned to roost later during summer which coincides with their breeding time. In the study sites, light intensity at the time of roosting was found to be significantly higher than the light intensity at the time of emergence. Similar findings have also been reported in Rook by Swingland (1976) where it was shown that Rooks departed from their roost at low light intensity and arrive at the roost at high light intensity. It may be possible that it is easier to find the roost site when it is brighter than under low-light conditions.

## Conclusion and Future Directions

Andrews and Naik (1970) and then Gaston (1977), carried out foundational work on the biology and social behaviour of Jungle Babbler, thereby presenting an excellent model system, to study sociality in a common backyard bird in the paleotropics. Yet, even three decades later, no serious attempt to follow on their seminal work was taken-up in the form of a rigorous scientific study of these cooperatively breeding passerine. In so, our study provides detailed information on their behavioural ecology with a quantified time-activity budget, diurnal and seasonal variation in activity patterns of the individual and social behaviour of Jungle Babblers. Several findings in Jungle Babblers with respect to time-activity budget were similar to social primates and were in contrast to what was found in similar studies on birds. This is likely due to the general paucity of time-activity budget studies on social birds and most similar studies in birds were carried out on solitary species, especially water birds. The similarity of our findings in Jungle Babblers to social primates highlights the importance of sociality in determining patterns of behaviour. Further, constraints on time allocation have been shown to impact aspects of social behaviour such as group size and it has been argued that activity budget needs to be included into models examining drivers of sociality (Pollard and Blumstein 2008). Studies on how animals allocate time to different activities according to their current status and surrounding environment will also be useful in providing valuable insight into how animals trade-off between different fitness enhancing behaviour and inform conservation policy making. Our study opens avenues for future studies examining the effect of ecological factors such as resource availability (abundant vs limited), habitat type (open versus closed), sociality (group size and composition) and predation pressure on the behaviour of Jungle Babblers in particular but of other social passerines in general.

## Acknowledgement

We thank Department of Forest and Wildlife Preservation, Government of Punjab, India for necessary permits (Permit No: 3625), Director NIPER for allowing us to carry out this study inside the campus and IISER Mohali for infrastructural support. We thank Sonam Chorol, Ranjit Singh and Gurtej for help with fieldwork. We are grateful to Anindya Sinha for his invaluable inputs on data analysis. We thank all members of BEL for suggestions and help with the work.

## Author Contributions

MJ conceived and designed the study and secured funding for the work. SDY carried out all the field work for data collection. SDY and MJ carried out all statistical analyses and wrote the manuscript.

## Data availability

The datasets generated are available as supplementary material

## Ethical approval

Jungle Babblers are listed in Schedule IV under the Indian Wildlife Protection Act (1972) and designated as ‘Least Concern’ by IUCN’s Red List of Threatened Species. This study was conducted with necessary permits (No. 3625) from Department of Forest and Wildlife Preservation, Government of Punjab, India, and with the approval of Institute Animal Ethical Committee (IISER/SAFE/PRT/2018/003), IISER Mohali, India. No animals were harmed or kept in captivity for this study.

