## Supplementary Material for "Temporal variation in the behaviour of a cooperatively breeding bird, Jungle Babbler (*Argya striata*) at diel and seasonal scale"

Table S1. Number of days and scans for every sampling hour

| Time | Days | Scans |
|---|---|---|
| 5-6 | 28 | 455 |
| 6-7 | 52 | 940 |
| 7-8 | 54 | 1130 |
| 8-9 | 36 | 645 |
| 9-10 | 40 | 825 |
| 10-11 | 37 | 780 |
| 11-12 | 50 | 905 |
| 12-13 | 49 | 975 |
| 13-14 | 43 | 925 |
| 14-15 | 44 | 740 |
| 15-16 | 47 | 930 |
| 16-17 | 42 | 885 |
| 17-18 | 62 | 1145 |
| 18-19 | 44 | 790 |
| 19-20 | 22 | 260 |

Table S2. Number of days and scans for each month

| Month | Days | Scans |
|---|---|---|
| October | 17 | 980 |
| November | 15 | 870 |

| | | |
|---|---|---|
| December | 15 | 930 |
| January | 9 | 385 |
| February | 14 | 495 |
| March | 16 | 865 |
| April | 16 | 1170 |
| May | 21 | 1585 |
| June | 17 | 1380 |
| July | 18 | 1280 |
| August | 21 | 1525 |
| September | 13 | 865 |
